# Rapid, large-scale, and effective detection of COVID-19 via non-adaptive testing

**DOI:** 10.1101/2020.04.06.028431

**Authors:** Matthias Täufer

## Abstract

Pooling of samples can increase lab capacity when using Polymerase chain reaction (PCR) to detect diseases such as COVID-19. However, pool testing is typically performed via an *adaptive testing strategy* which requires a feedback loop in the lab and at least two PCR runs to confirm positive results. This can cost precious time. We discuss a non-adaptive testing method where each sample is distributed in a prescribed manner over several pools, and which yields reliable results after one round of testing. More precisely, assuming knowledge about the overall incidence rate, we calculate explicit error bounds on the number of false positives which scale favourably with pool size and sample multiplicity. This allows for hugely streamlined PCR testing and cuts in detection times for a large-scale testing scenario. A viable consequence of this method could be real-time screening of entire communities, frontline healthcare workers and international flight passengers, for example, using the PCR machines currently in operation.

## 1. Introduction

One key to containing and mitigating the COVID-19 pandemic is suggested to be rapid testing on a massive scale [HZW+20, SBY]. It would be beneficial to develop the ability to routinely, and in particular rapidly, test groups such as frontline healthcare workers, police officers, and international travellers. Testing for SARS-CoV-2 is currently performed via the polymerase chain reaction (PCR) on nasopharyngeal swabs [TTY+20]. Typically, the population size significantly exceeds the capacity for testing, with the number of available PCR machines and reagents an important bottleneck in this process.

There are two basic approaches to PCR testing in populations: 1. individual tests, where every single sample is examined, and 2. pooled tests where larger sets of samples are mixed and tested en bloc. Pooled testing was pioneered by Dorfman in 1943 [Dor43] in the context of blood tests and led to a host of research activity, both on the lab side as well as the theoretical side [AJS19, DH99, DH06]. If the disease is rare in the population, pooled testing may be advisable. In this case it can assist in optimizing precious testing capacity since most individual results would be negative. Pooling relies on the fact that the PCR is reasonably reliable under the combination of samples: the preprint [YAST+20] suggests that a detection of SARS-CoV-2 in pools of size 32 and possibly 64 is feasible.

While a classic pooling strategy has the advantage that less overall PCR tests are required, there are disadvantages in terms of lab organisation and – more crucially – time: pooling only indicates whether a pool contains at least one infected individual. If samples are tested in pools of size *n* and the incidence *ρ* is small (more precisely, if *ρ · n* is small) then a number of samples will be in pools that are tested positive and hence undergo a second round of testing. In other words, pooled testing with individual verification of positive pools is an *adaptive testing strategy*, the lab organisation for which is a labour, management, and resource intensive process. It has several drawbacks, since it requires keeping multiple lab samples and re-running of the time-intensive PCR process. The lab feedback loop makes the entire workflow more susceptible to delays (see Figure 2). This may result in delays in individual results – a particular problem when the objective is to rapidly identify infected individuals, who may infect others while waiting for the test outcome. Furthermore, since the number of samples undergoing a second round of testing is an unknown quantity, some reserve capacity is required to prevent further delays. This makes it more challenging for the lab to operate near its maximal capacity.

**Figure 1:**
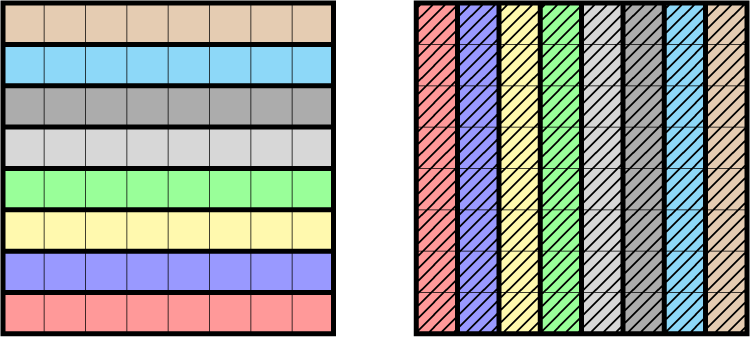
Pooling along rows and columns to arrange *N* = 64 samples into 16 pools of size 8 to form a (64, 8, 2)-multipool. Different background patterns and colours represent different pools.

**Figure 2:**
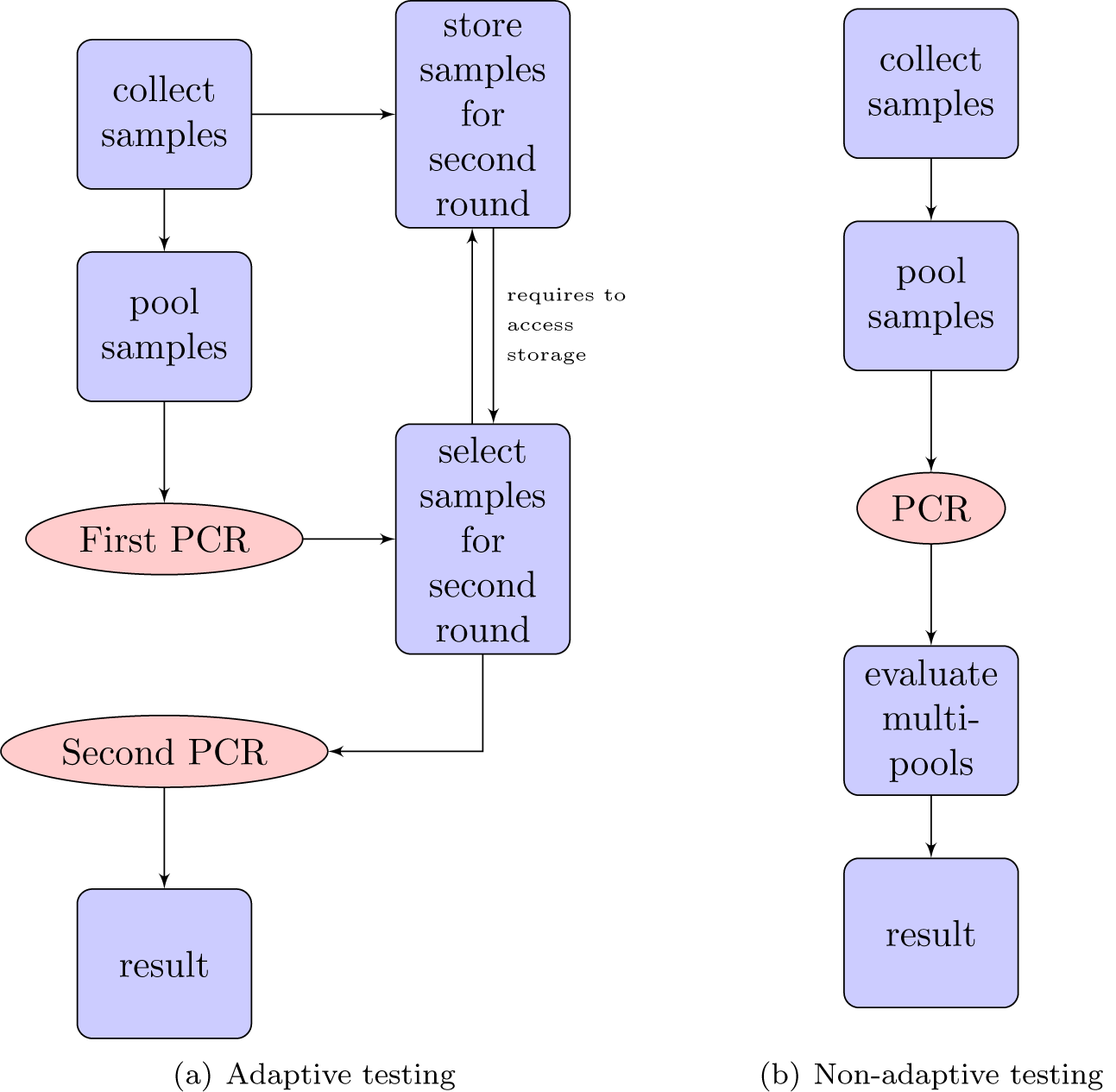
Comparison of the work flow in adaptive testing and non-adaptive testing. In the adaptive setting in Figure (a), two of the time-expensive PCR steps (in red) are required. Furthermore, the interplay of data interpretation after the first PCR and the sample storage management introduces another possible bottleneck. In the non-adaptive case, shown in Figure (b), the work flow is completely linear.

In the theoretical research on testing strategies the distinction is made between *adaptive testing*, for example when all samples in a positive pool undergo a second round of testing, and *non-adaptive* strategies, where all tests can be run simultaneously [DH99]. Testing every sample individually can be considered as a trivial non-adaptive strategy, but there exist non-adaptive strategies which combine the benefit of pooling with the advantages of non-adaptive testing.

In this note, we propose a non-adaptive pooling strategy for rapid and large-scale screening for SARS-CoV-2 or other scenarios where detection time is critical. This allows for significant streamlining of the testing process and reductions in detection time. Firstly because only one round of PCR is required, and secondly because it eliminates actions in the lab workflow that require input from results determined in the lab, i.e. the testing infrastructure can be organized completely linearly, cf. Figure 2 for an illustration. The strategy will systematically overestimate the number of positives, but we can provide error bounds on the number of false positives which scale favourably with large numbers and will be small in realistic scenarios.

## 2. Definition of the non-adaptive testing strategy: multipools

Our testing strategy is as follows: every individual’s sample is broken up into *k* samples and distributed over *k* different pools of size *n* such that no two individuals share more than one pool. An individual is considered as tested positive if all the pools in which its sample has been given are tested positive or – in our case equivalently – an item is considered as tested negative if it appears in at least one negative pool. This decoding algorithm is also known as COMP (Combinatorial Orthogonal Matching Pursuit), an algorithm easily implementable in practice with low run-time and storage [JAS19].

Let us make our definition more formal:

### Definition 1 (Multipools).

*Let a* population *(X*_1_, *…, X*_*N*_ *) of size N, a* pool size *n, and a* multiplicity *k be given, and assume that Nk is a multiple of n. We call a collection of subsets/pools of* {*X*_1_, *…, X*_*N*_} *an* (*N, n, k*)-multipool, *or briefly* multipool, *if all of the following three conditions hold:*

*(M1) Every pool consists of exactly n elements*.

*(M2) Every sample X*_*i*_ *is contained in exactly k pools*.

*(M3) For any two different samples X*_*i*_, *X*_*j*_ *there exists at most one pool which contains both X*_*i*_ *and X*_*j*_.

In the context of non-adaptive testing, designs as in Definition 1 are called (*k* − 1)*-disjunct matrices* and it is known that such matrices correctly identify up to *k* infected samples [Maz12]. However, we will be interested in scenarios where the number of infected samples can exceed the multiplicity *k*. If *N* = *n*^2^ and *k* = 2 the construction of an (*N, n*, 2)-multipool is quite straightforward, see Figure 1: arrange the *N* samples in a rectangular grid and then pool along every row and column, cf. [SSW+16, FFLH, ZDF+14]. However, as we shall see below, *k* = 2 is in many realistic scenarios insufficient for the desired precision.

Some recent contributions [FFLH, MNB+20] propose to arrange samples in a (3 or higher dimensional) hypercube and to pool along all hyperplanes. This makes every individual sample appear in three or more pools, but it is *not* a multipool in the sense of Definition 1 above, since in dimension three and higher, any two hyperplanes will intersect in more than one point, in violation of Property (M3). This creates unnecessary correlations between different pools and impairs performance. If *k* = 3, systems as in Definition 1 are also called *Steiner triples* and have been recently used in non-adaptive group testing for SARS-CoV-2 [GAR+20]. A flexible way to construct multipools of various multiplicities *k* is given by the Shifted Transversal Design [TM06, EGN+15] which we explain in Section 4.

## 3. Controlling the number of false positives

We always assume that the incidence *ρ* of the disease is small compared to the inverse pool size 1*/n*. This is a reasonable requirement, also in classical pooling strategies (a *ρn* portion of samples will have to undergo second testing, thus a large *ρn* would attenuate the benefit of pooling).

Assuming perfect performance of the PCR, also under pooling (see Section 6 on how to deal with uncertainty here), multipooling will identify all infected individuals, since all their pools will be positive. However, a sample might falsely be declared positive if all pools in which it is contained happen to contain an infected sample.

The expected portion of false positives in a multipool strategy is

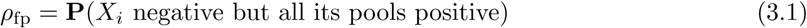

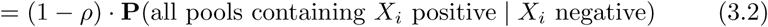

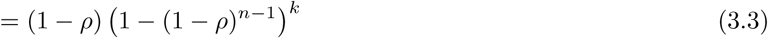

Here, the third identity crucially uses the property (M3) which guarantees independence between the poolmates in the different pools of a sample. By Bayes’ rule, the probability to actually be negative when tested positive by the multipool (i.e. the portion of subjects falsely declared positive among all subjects declared positive) is

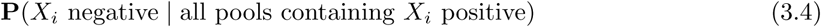

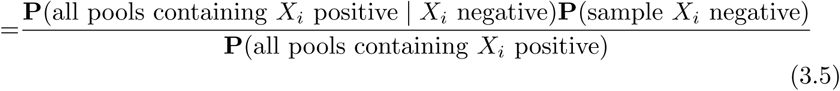

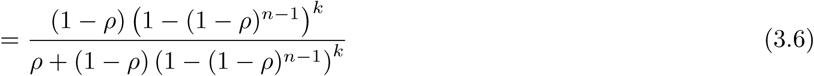

Let us calculate for which *k* the probability of a positive test result being a false positive does not exceed *ϵ*_fp_ *>* 0:

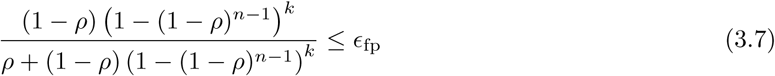

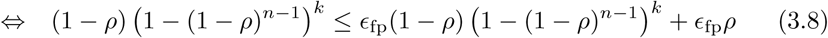

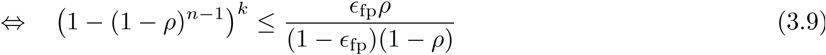

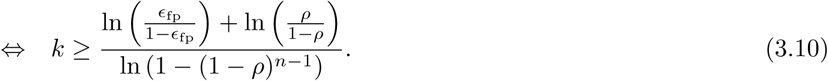

This provides a lower bound on the necessary multiplicity *k* in terms of the sample size *n*, the knowledge on the incidence *ρ*, and the acceptable portion *ϵ*_fp_ of false positive results among all positives. Assuming *ϵ*_fp_ *<* 1 and 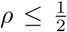 (which are both reasonable assumptions, recall that *ρn* is small), the lower bound in (3.10) is monotone increasing in *ρ*. Hence, if the exact incidence is unknown but we have an upper bound on it, we can work with the largest/worst case *ρ*. Let us summarize these findings in the following

### Theorem 1.

*Let the incidence be at most* 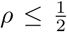, *and let* 0 *< ϵ*_fp_ *<* 1. *If*

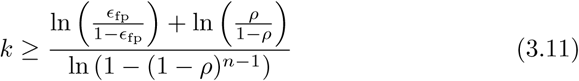

*then in any multipooling strategy with pool size n and multiplicity k, the probability of a positive test being a false positive does not exceed ϵ*_fp_.

The number of tests required in a multipool strategy is *Nk/n*, an improvement compared to individual testing by a factor *n/k*. A key observation is that the lower bound on *k* in Inequality (3.11) scales favourably with large multiplicities *n*. Indeed, recall that in an adaptive pooling strategy one wants on the one hand large pool sizes *n*, but on the other hand *nρ* should be small. It is therefore reasonable to have *n* proportional to the inverse of *ρ*, i.e. *nρ* ≈ *C*. Using that 1 −*ρ* ≈ 1 and 1 − (1 −*ρ*)^*n*−1^ ≈ (*n* − 1)*ρ* ≈ *nρ*, the lower bound in (3.11) behaves approximately as

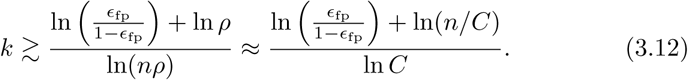

that is *k* grows only logarithmically with the pool size *n*. An analogous analysis shows that *k* also grows logarithmically with the inverse of *ϵ*_fp_ when the error probability *ϵ*_fp_ is sent to zero. We compare the theoretical values found in the lead-up to Theorem 1 to simulated values in Figure 3

**Figure 3:**
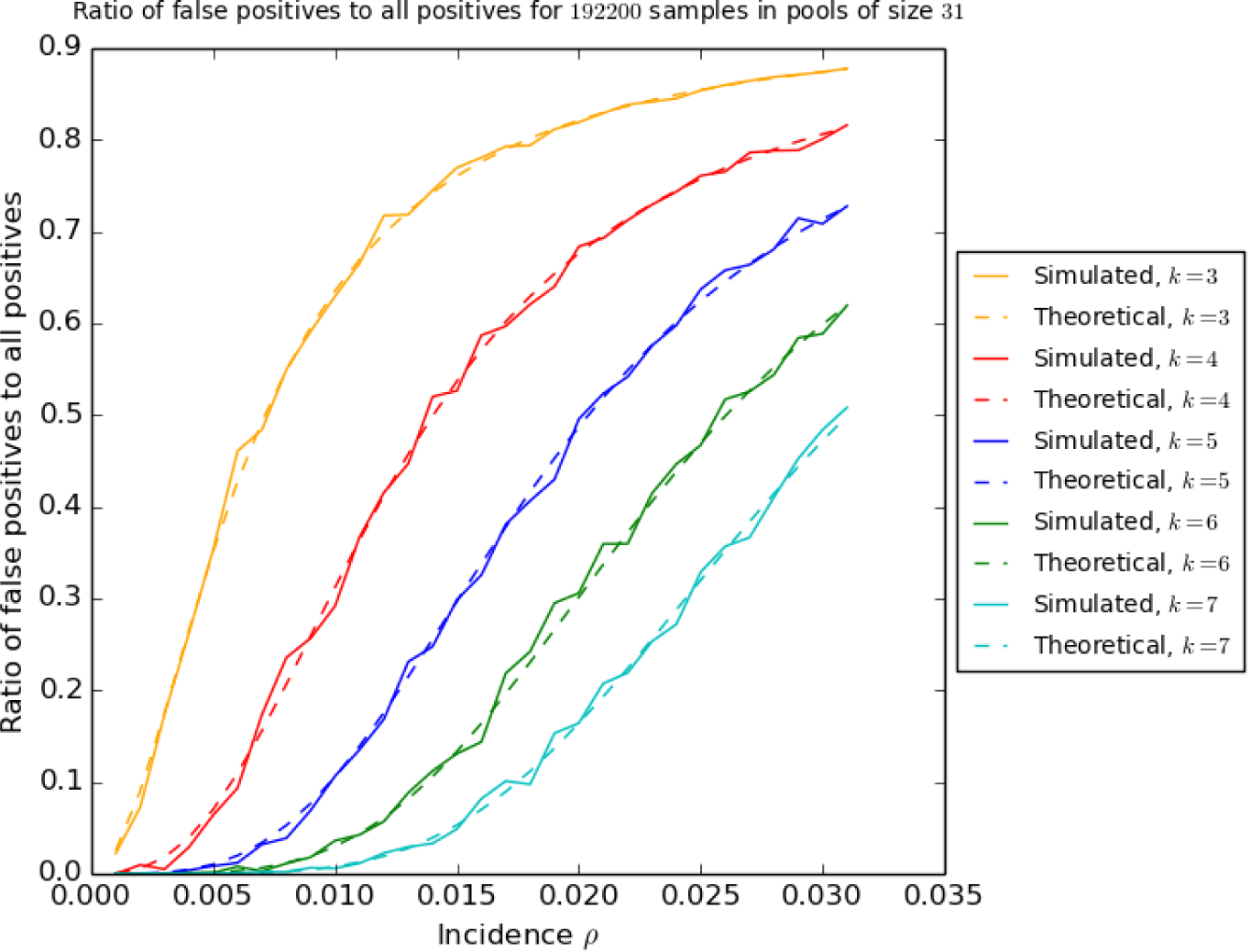
Comparison of the ratio of false positives to positive results in simulations on synthetic data for 200 931 samples with different incidences *ρ* at pool size *n* = 31 and sampling strategies with multiplicities *k* ∈ {4, 5, 6, 7}, and the theoretical value calculated in the lead-up to Theorem 1. The code for the simulation can be found in [Täu20].

## 4. Generating multipools

The question for which combiniations (*N, n, k*) a multipool exists seems to be in general a non-trivial combinatorial problem. We focus here on the case when *N* = *n*^2^ and on constructions based on the Shifted Transversal Design [TM06].

It is useful to imagine all *N* samples arranged in an *n* × *n*-square and label samples by their *x* and *y*-coordinate, i.e. denoting the sample at position 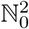 by *X*_*ij*_, where we define the the sample in the lower left (south-west) corner to be *X*_00_. For multiplicity *k* = 2, a (*N, n, k*)-multipool can be constructed by pooling along rows and columns, as in Figure 1.

Unfortunately, for reasonable parameter choices, a multiplicity of *k* = 2 turns out to lead to large false positive rates: For instance, arranging *N* = 64 samples from a population with incidence *ρ* = 0.01 in a rectangular grid and pooling along all rows and columns (in our notation this is an (64, 8, 2)-multipool), Identity (3.6) will imply that on average 31.4% of positive results will actually be false positives. To improve on this and pass to multiplicity *k* = 3, one can sample along diagonals, where the diagonals are continued periodically, see Figure 4. This works for any pool size *n* ≥ 2 and leads to

**Figure 4:**
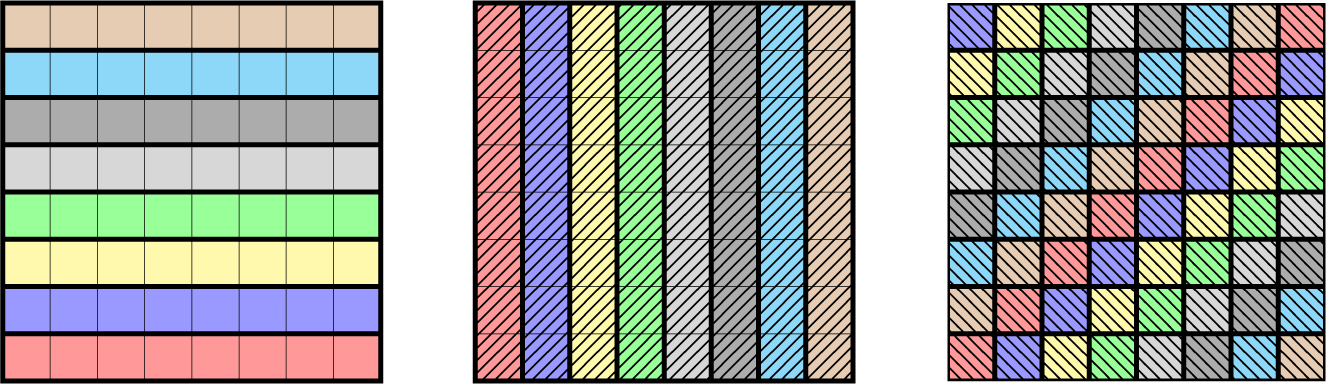
Pooling along rows, columns, and periodically continued diagonals to arrange *N* = 64 samples into 24 pools of size 8 to form a (64, 8, 3)-multipool. Different background patterns and colours represent different pools.

### Theorem 2.

*Let N* = *n*^2^ *and n* ≥ 2. *Then there exists an* (*N, n*, 3)*-multipool, obtained by sampling along rows, columns, and all periodically continued south-west-to-north-east diagonals*.

In the situation of *N* = 64 and *n* = 8, this allows for the construction of a (64, 8, 3)-multipool in which, by (3.6), the probability of a positive result being erroneous is reduced to 3.01%. In such a scenario, one would test 64 individuals with 24 tests, a compression by a factor 0.375. A higher compression rate would require larger pool sizes *n*. Since the lower bound (3.11) on *k* in Theorem 1 is monotonous in *n*, this will in turn also require to higher multiplicities *k* in order to achieve comparable false positive error probabilities. To pass to *k* = 4, one might now be tempted to pool along the other (north-west-to-south-east) diagonals, but this is not going to yield a multipool in general, see for instance Figure 5 where, in the case *n* = 8, two diagonals intersect in more than one point, in violation of Property (M3) in Definition 1.

This is due to the fact that *n* = 8 has non-trivial divisors, i.e. it is not a prime number. South-west-to-north-east diagonals are of the form

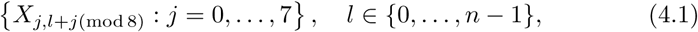

and north-west-to-south-east diagonals

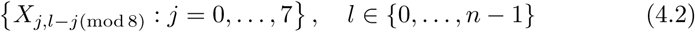

were, (mod *n*) means that we use arithmetic modulo *n*, that is as soon as we exceed *n* − 1, we start counting from 0 again. These diagonals are lines of slope +1 and −1, respectively, and the difference of these slopes is 2, which divides 8. Since intersections of two such lines are given by solutions to the equation

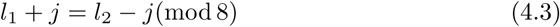

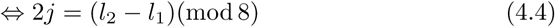

there can be more than one *j* solving (4.4): Indeed, if some *j*_0_ ∈ {0, *…*, 7} solves (4.4), then *j′*:= *j* + 4(mod 8) is a solution as well, since 2*j*^*′*^= 2*j*(mod 8).

More generally, it is well-known that for *m* ∈ {1, *…, n* − 1} and *j* ∈ {0, *…, n* − 1}, the equation

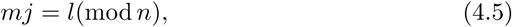

has a unique solution *j* if and only if the greatest common divisor of *m* and *n* is 1. Since this must hold for all *m* ∈ {1, *…, n* − 1}, *n* must be a *prime number*. In this case, the integers modulo *n* form an algebraic structure called a *field*, in which every non-zero element has a well-defined multiplicative inverse. For prime *n*, the unique solution of (4.5) is therefore given by *j* = *m*^−1^*l*, where *m*^−1^ denotes the multiplicative inverse of *m* in arithmetic modulo *n*.

This suggests to use a prime pool size *n* and sample along lines of different slopes, that is to use pools of the form

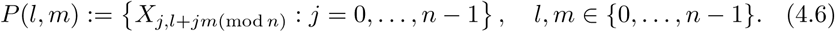

We can add one more type of pool by sampling along all vertical lines (their slope can be considered as “infinity”) which we denote by

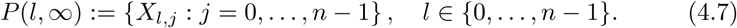

Such ensembles of pools are sketched in Figure 6 for the case *n* = 5.

This construction is also referred to as the *Shifted Transversal Design* in [TM06]. We summarise our findings in the following

**Figure 5:**
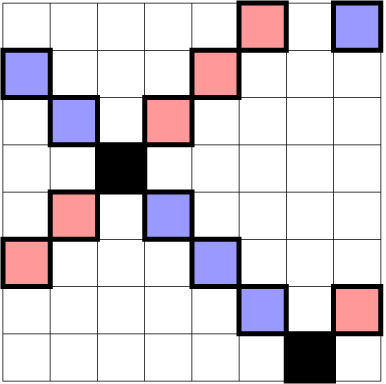
The two diagonals (red and blue) intersect in two points (black). They cannot both be used as pools in a multipool.

**Figure 6:**
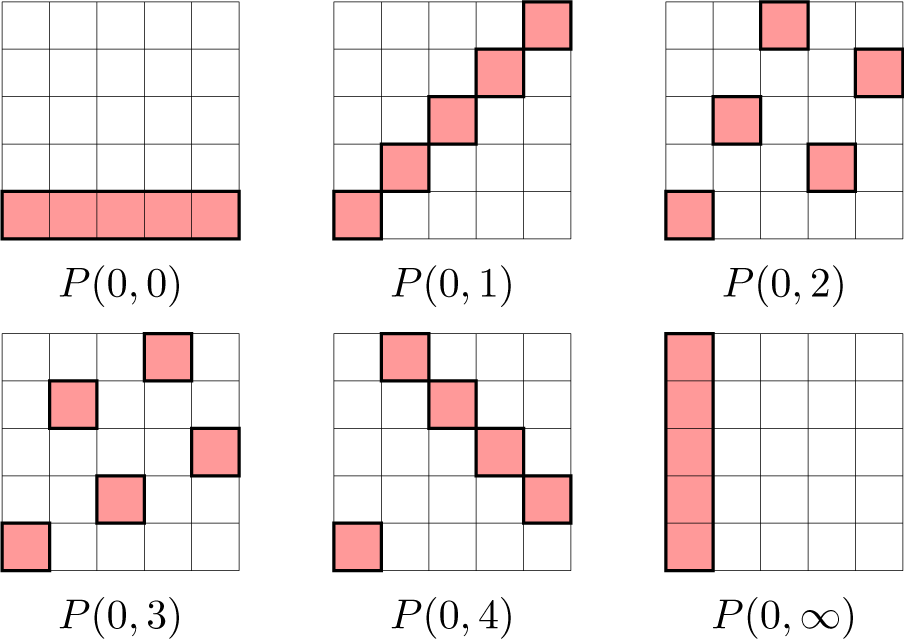
Pools of different slopes as in Theorem 3 for *n* = 5.

### Theorem 3.

*Let n be a prime number and let N* = *n*^2^. *Then, there exists a* (*N, n, k*)*-multipool for k* = (*n* + 1), *and consequently also for every smaller k. This multipool is given by pooling along all sloped lines, that is:*

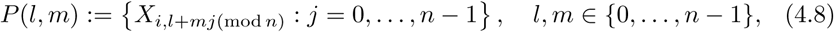

*and pooling along all columns (or lines of slope infinity), that is*

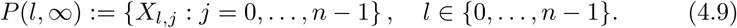

Figure 6 contains an illustration of elements of such a multipool in the case *n* = 5 with multiplicity *k* = 6. Theorem 3 allows for multiplicities up to *k* = *n* + 1, but in practice, one will want to work with much lower multiplicities *k* since a high multiplicity would require many tests and defeat the purpose of pooling. From a practical perspective it seems reasonable to generate large pools by a sequence of unions of two equally diluted pools. This leads to pool sizes which are a power of 2, certainly not a prime number (except for 2 itself). One approach to accomodate for that would be population sizes *N* = *n*^2^ where *n* is a prime just below a power of 2, e.g. *n* = 31, which is just below 32 or *n* = 61 which is just below 64. Then pools of size *n* can be mixed by adding a small number of negative dummy samples and proceeding as if *n* was a power of 2.

## 5. Examples and scenarios

Let us sketch some concrete examples where the pool sizes are a prime number and where the multipooling strategy might be useful:

*N = 961, ρ* ≤ 1%, *n* = 31

Let the population size be *N* = 31^2^ = 961. This could for instance be the number of employees in a company or passengers which depart from an international airport within a certain time window. Let the incidence rate *ρ* be no more than 1.0% and let us work with a pool size *n* = 31. Since *n* is prime, Theorem 3 allows to construct (961, 31, *k*)-multipools for any *k* ≤ 32 and Theorem 1 allows to bound the probability of a positive test being erroneous for different multiplicities *k* as in Table 1. Accepting for instance a false positive probability of 3% requires 6*N/n* = 186 PCR tests, 19.4% of what would be required in individual testing. Let us emphasize again here that this means that 3% among the *results flagged as positive* will be false positives, not 3% of the overall test results.

**Table 1:** Probability of a positive result being a false positive and the compression *k/n* compared to individual testing for pool size *n* = 31, incidence *ρ* ≤ 0.01 and different multiplicities *k*.

| $k$ | $\epsilon_{\text{fp}}$ | $k/n$ |
| --- | --- | --- |
| 4 | 0.32 | 0.129 |
| 5 | 0.11 | 0.161 |
| 6 | 0.03 | 0.194 |
| 7 | 0.008 | 0.226 |

*N = 3721, ρ* ≤ 0.1%, *n* = 61

The multipool method scales well with larger numbers. Let the population size be *N* = 61^2^ = 3721 and the pool size *n* = 61, which is of the order of pools being used for the PCR today [YAST+20]. Let furthermore be the incidence rate be no larger than 0.1%, a realistic upper bound for the prevalence of SARS-CoV-2 in many countries [fNS20]. Since *n* = 61 is prime, Theorem 3 allows to construct (3721, 61, *k*)-multipools for any *k* ≤ 62 and the error bounds in Theorem 1 lead to Table 2. If we choose *k* = 4 and accept *ϵ*_fp_ = 1.2% as the probability for positive results being false positives, we need 4*N/n* = 244 tests in order to fast and efficiently test 3721 individuals, that is 6.6% of what would be needed with individual testing.

**Table 2:** Probability of a positive result being a false positive and the compression *k/n* compared to individual testing for pool size *n* = 31, incidence *ρ* ≤ 0.01 and different multiplicities *k*.

| $k$ | $\epsilon_{\text{fp}}$ | $k/n$ |
| --- | --- | --- |
| 3 | 0.17 | 0.049 |
| 4 | 0.012 | 0.066 |
| 5 | 0.0007 | 0.082 |

## 6. Discussion and possible extensions

The non-adaptive multi-pooling strategy provides a streamlined and efficient organisation of the testing process and cuts in detection time. This significant benefit comes with potential reductions in accuracy compared with adaptive testing, but this false positive rate can be tightly controlled and tailored to suit the circumstance. The false positive probability *ϵ*_fp_ deemed an acceptable cost for the increased testing efficiencies may depend on, for example, the infection characteristics, the government policy and resource levels.

A small modification of our strategy might furthermore allow for an improvement of the false negative rate – even compared to usual adaptive pool testing strategies: even though commonly used, pooling samples can potentially dilute samples close to the identification threshold of the PCR and increase the probaility of false negatives. The recent preprint [YAST+20] estimates a false negative rate of 10% when detecting SARS-CoV-2 in pools of size 32. One can reduce this type of false negative in our strategy by declaring all samples which are *in at least k* − 1 *positive pools* as tested positive.

This strategy is known as the “Noisy COMP” (NCOMP) decoding algorithm [CCJS11, CJSA14] where an item is declared infected if more than a certain portion of its pools test positive. This will on the one hand lower the probability of false negatives, but more importantly it will only mildly affect the false positive rate. This could be seen by adding a next-order term in the error analysis performed leading up to Theorem 1. For a sound analysis, knowledge on the false positive rate gained through experiments would be required, but the general message that the necessary multiplicity *k* will grow slowly with large *n* and small *ϵ*_fp_ remains.

Let us finally note that the basic idea is close to compressed sensing and sparse recovery [CT06, FR13]. While in our situation the output space consists of {0, 1} -vectors, which make the mathematics we use rather elementary, there also seem to be applications of the PCR where quantitative measurements are taken and where compressive sensing techniques might be applied. A very recent approach in this direction is *Tapestry pooling* [GRK+20, GAR+20] which takes quantitative data from PCR measurements and uses methods from compressed sensing to decode. In the scenario of testing *N* = 961 samples in pools of size *n* = 31 discussed in Section 5, this approach suggests reasonable results at multiplicity *k* = 3, a higher compression rate than in our approach. However we emphasise that the (experimental) error analysis performed in the context of Tapestry pooling focuses on fixed numbers of infected samples and is therefore in a slightly different spirit than our approach which is based on the prevalence of the disease in the population.

## Acknowledgements

The author thanks Christoph Schumacher for numerous helpful discussions and comments. Comments by Emma Lawrance, Albrecht Seelmann and Sasha Sodin are also gratefully acknowledged.

